# Skeletal Muscle Mitochondrial Dysfunction Mediated by *Pseudomonas aeruginosa* Quorum Sensing Transcription Factor MvfR: Reversing Effects with Anti-MvfR and Mitochondrial-Targeted Compounds

**DOI:** 10.1101/2024.05.03.592480

**Authors:** Shifu Aggarwal, Vijay Singh, Arijit Chakraborty, Sujin Cha, Alexandra Dimitriou, Claire de Crescenzo, Olivia Izikson, Lucy Yu, Roberto Plebani, A. Aria Tzika, Laurence G Rahme

## Abstract

Sepsis and chronic infections with *Pseudomonas aeruginosa,* a leading “ESKAPE” bacterial pathogen, are associated with increased morbidity and mortality and skeletal muscle atrophy. The actions of this pathogen on skeletal muscle remain poorly understood. In skeletal muscle, mitochondria serve as a crucial energy source, which may be perturbed by infection. Here, using the well-established backburn and infection model of murine *P. aeruginosa* infection, we deciphered the systemic impact of the quorum sensing (QS) transcription factor MvfR by interrogating five days post-infection its effect on mitochondrial-related functions in the gastrocnemius skeletal muscle and the outcome of the pharmacological inhibition of MvfR function and that of the mitochondrial-targeted peptide, Szeto-Schiller 31 (SS-31). Our findings show that the MvfR perturbs ATP generation, oxidative phosphorylation (OXPHOS), and antioxidant response, elevates the production of reactive oxygen species, and promotes oxidative damage of mitochondrial DNA in the gastrocnemius muscle of infected mice. These impairments in mitochondrial-related functions were corroborated by the alteration of key mitochondrial proteins involved in electron transport, mitochondrial biogenesis, dynamics and quality control, and mitochondrial uncoupling. Pharmacological inhibition of MvfR using the potent anti-MvfR lead, D88, we developed, or the mitochondrial-targeted peptide SS-31 rescued the MvfR- mediated alterations observed in mice infected with the wild-type strain PA14. Our study provides insights into the actions of MvfR in orchestrating mitochondrial dysfunction in the skeletal murine muscle, and it presents novel therapeutic approaches for optimizing clinical outcomes in affected patients.

## INTRODUCTION

Pathogens manipulate mitochondrial dynamics and functions to enhance their survival or evade host immunity (1). In skeletal muscle, mitochondria play a vital role as an energy powerhouse, essential for maintaining cellular homeostasis (1, 2). One of the primary roles of mitochondria is to generate adenosine triphosphate (ATP) through oxidative phosphorylation (OXPHOS) to fuel cellular functions (3). The number of mitochondrial copies per cell is related to the cell’s energy demands, and the mitochondrial DNA (mtDNA) copy number varies depending on the cell’s energy needs and oxidative stress levels (4–6). These important organelles establish a dynamic network by constantly undergoing fusion and fission to remove aging or damaged mitochondria in response to cellular stress and maintain mitochondrial functions (7–9). They adapt to diverse physiological stresses, including infection, by regulating many cellular activities and host responses during bacterial infections, underscoring their pivotal role in maintaining homeostasis during infection (1). Numerous bacteria are recognized for their ability to restructure host cell metabolism to improve their intracellular survival, with most decelerating the TCA cycle and promoting aerobic glycolysis (1). Alterations in metabolism may reduce respiration and increase reactive oxygen species (ROS) generation that damages mitochondrial DNA (mtDNA), proteins, and lipids (7, 10, 11). Bioenergetic failure or any defects or abnormalities in mitochondria that can result from ATP depletion, excessive ROS levels (12) and as part of autophagy, mitophagy, and apoptosis (13–15) can affect skeletal musculature and result in multisystem diseases (16).

Sepsis and severe thermal injuries may lead to skeletal muscle loss. Similarly, cystic fibrosis (CF) patients also experience skeletal muscle atrophy and dysfunction due to a lack of the CF transmembrane conductance regulator (CFTR) (17). Critically ill patients, like cystic fibrosis and patients with burns, are highly prone to *P. aeruginosa* infections, augmenting the adverse effects of these pathologies on skeletal muscle (18–20). Especially because skeletal muscle constitutes 40% of total body weight and plays numerous vital roles in the human body, including movement, posture maintenance, breathing facilitation, and safeguarding internal organs (21).In previous studies, we have shown that the *P. aeruginosa* quorum sensing (QS) transcription factor MvfR (<u>M</u>ultiple <u>v</u>irulence factor <u>R</u>egulator) (22, 23), also known as PqsR, is a central QS regulator that controls the expression of multiple virulence genes and the synthesis of many small signalling molecules in this pathogen. MvfR is required for *P. aeruginosa* full virulence (24–26). It directly regulates the expression of more than 30 genes, including the *pqsABCDE* and *phnAB* operons that catalyze the biosynthesis of ∼60 distinct low-molecular-weight compounds (23, 25, 27, 28) part of which are the 4-hydroxy-2 alkylquinolines (HAQs) that include the signalling molecules and MvfR inducers 4-hydroxy-2-heptylquinoline (HHQ), 3,4-dihydroxy-2-heptylquinoline (PQS), as well as 2-n-heptyl-4-hydroxyquinoline-N-oxide (HQNO) and the non-HAQ molecule 2-aminoacetophenone (2-AA) that promotes persistent infections (26, 29–33). In a series of studies, we have shown the impact of 2-AA’s on immune cells and skeletal muscle functions(24, 26, 29, 30, 34, 35). In non-infection studies, we have shown that this MvfR-regulated small signalling molecule impacts mitochondrial-related functions in skeletal muscle decreases the ATP synthesis rate in the gastrocnemius muscle of live animals (34, 36).

Given that MvfR is required for full virulence *in vivo* and controls multiple virulence functions, including the synthesis of 2-AA, in this study, we interrogated the impact of this *P. aeruginosa* master QS transcription regulator in mitochondrial homeostasis and related functions during infection. Moreover, we examined whether pharmacological inhibition of MvfR function using the N-Aryl Malonamides (NAM) compound D88 we developed (37) could mitigate MvfR mediated mitochondrial and related function derangements during *P. aeruginosa* infection. We have demonstrated that MvfR activity can be inhibited *in vivo* by the NAM compound D88 (37). D88 is highly efficacious in mitigating MvfR virulence and inhibiting the synthesis of HAQs and 2-AA by binding to the same ligand-binding domain (LBD) hydrophobic pocket as its inducers HHQ and PQS (24, 25, 37). Using this NAM compound in monotherapy, we have shown that it protects murine intestinal barrier function, abolishes the synthesis of the small signalling under MvfR control, ameliorates bacterial dissemination, and lowers inflammatory cytokines (37). Furthermore, to also determine whether mitochondrial dysfunction mediated by *P. aeruginosa* could be counteracted by a compound targeting mitochondria rather than the pathogen, we used the Szeto–Schiller mitochondrial-targeted peptide 31 (SS-31), an amphipathic tetrapeptide shown to protect cells against induced oxidative stress, reducing intracellular ROS, and maintaining membrane potential (ΔΨm) (38–40). Our findings provide evidence and insights into the role of this QS regulator in metabolic perturbances linked to mitochondrial functions in skeletal muscle in an infection setting *in vivo* and offer therapeutic approaches through the pharmacologic inhibition of MvfR by a highly efficacious anti-MvfR compound we developed and the use of the mitochondria-targeted peptide, SS-31.

## RESULTS

### MvfR perturbs ATP generation, oxidative phosphorylation, and ROS production in gastrocnemius murine muscle

We have shown that injecting mice with the MvfR-regulated molecule 2-AA decreases the ATP synthesis rate and the twitch tension of the tibialis and gastrocnemius muscle (34). These data prompted us to interrogate the impact of the MvfR on the ATP levels and mitochondrial-related proteins in the gastrocnemius muscle 5 days post-infection in mice infected with the wild-type *P. aeruginosa* strain PA14 and compare them to mice infected with the isogenic Δ*mvfR* mutant strain. The sham mice group were basal line control (Figure 1A).

**Figure 1:**
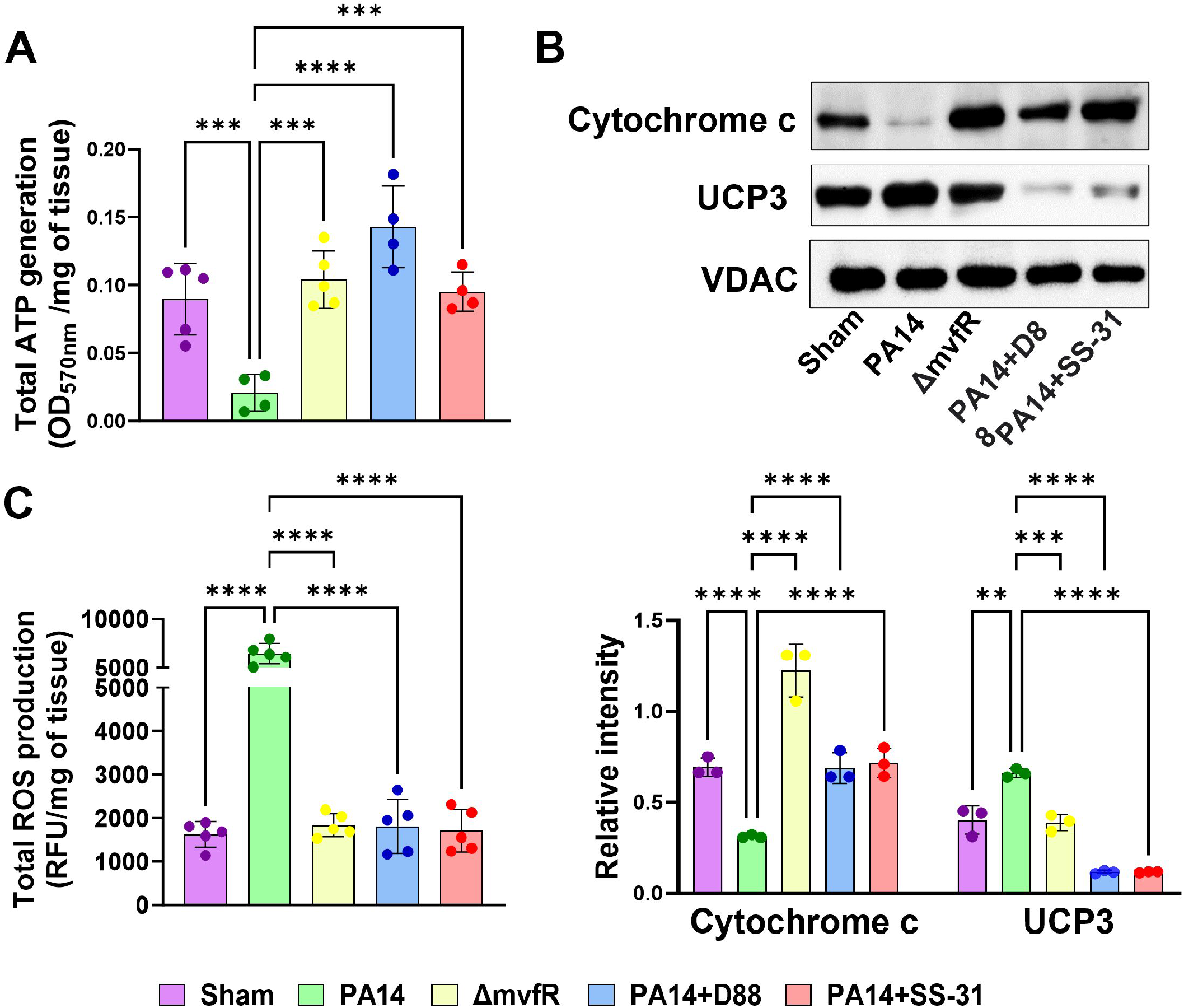
MvfR decreases ATP levels by impacting mitochondrial cytochrome c protein and increases the uncoupling protein UCP3 in gastrocnemius muscle. These alterations are reinstated by the pharmacological inhibition of MvfR or administration of SS-31. (A) Total ATP generation calculated as OD_570nm_ per milligram tissue (B) Representative Western blot images of cytochrome c and UCP3 using mitochondrial lysates from the gastrocnemius muscle of sham and infected mice with wild-type strain PA14 or Δ*mvfR* or PA14-infected and treated with the anti-MvfR compound D88, or the mitochondrial-targeted SS-31 peptide. Bar charts (bottom) depicting the relative signal intensity versus total protein loading amount for skeletal tissues of three mice (n=3). (C) Total ROS production depicted as relative fluorescence unit per gram of tissue (RFU/mg of tissue) in the gastrocnemius muscle of mice infected with PA14 strain (green), Δ*mvfR* strain (yellow), anti-mvfR compound D88 (blue) and mitochondrial-targeted peptide SS-31 (red), when added separately to the PA14 infected mice. Sham (purple) representing the burn group was used as a baseline control. Each dot represents muscle tissue from five mice (n=5). Error bars represent the standard deviation (SD) from the average of five mice. The statistical difference between each protein level is shown where *** p<0.001 and **** p < 0.0001 indicate significance. VDAC was used as a loading control. One-way ANOVA followed by Tukey’s post hoc test was applied.

Our findings show a significant decrease in the total ATP generation with PA14 infection compared to skeletal muscle samples from the Δ*mvfR-*infected mice. ATP generation in mitochondria involves the coordinated action of various electron transport chain proteins and oxidative phosphorylation proteins. Cytochrome c is a multifunctional enzyme, acting as an essential electron transport carrier, and maintains the mitochondrial transmembrane potential essential for ATP generation (41, 42). Hence, we investigated whether the MvfR function impacts this electron transport chain protein localized to the mitochondrial intermembrane space. Western blotting analysis of cytochrome c (Figure 1B) protein from the mitochondrial fraction of the skeletal muscle of infected mice showed that PA14 infection significantly decreases the levels of this protein compared to the Δ*mvfR* infected mice that exhibited even higher levels than the sham control baseline group. This finding corroborates the ATP result and supports the contributory role of MvfR in impairing the mitochondrial transmembrane potential and decreasing energy metabolism, which is associated with the reduced generation of ATP.

Since MvfR reduces the cytochrome c protein levels and this protein can operate as a reactive oxygen species (ROS) scavenger (43, 44), we measured ROS levels in the skeletal muscle of infected mice (Figure 1C). In agreement with the above findings, the assessment of the total ROS production in the gastrocnemius muscle showed that in PA14-infected mice, its production is dramatically increased as opposed to the levels in the tissues of the Δ*mvfR* infected mice, which were found to be similar to the baseline levels of the sham control group (Figure 1C). These results also prompted us to assess the levels of uncoupling protein 3 (UCP3). Uncoupling proteins regulate mitochondrial membrane potential and key steps in cellular bioenergetics (45). Specifically, UCP3 plays an active role in ROS production in skeletal muscle, and thus, we assessed its levels in the mitochondrial fraction of skeletal muscle protein extracts from infected mice. Figure 1B also shows that in corroboration with the above findings, PA14 infection led to increased mitochondrial UCP3 protein levels in skeletal muscle, thus signifying the impact of MvfR on oxidative metabolism and the oxidation status of the tissue.

### The anti-MvfR compound D88 and the mitochondrial-targeted peptide SS-31 rescue the metabolic alterations promoted by MvfR

Subsequently, we aimed to determine the therapeutic efficacy of the anti-MvfR compound D88 and the mt-targeted peptide SS-31 in the alterations observed in the skeletal muscle of infected mice. Both compounds reinstated the ATP generation, ROS production, and the cytochrome c protein levels in the skeletal muscle of the PA14-infected animals to similar levels observed in Δ*mvfR-infected* or sham control group mice (Figure 1A-C). The UCP3 expression in PA14-infected animals was also rescued by the treatment of D88 or SS-31, with both compounds reducing the levels of this uncoupling protein slightly below the sham control basal or *mvfR* triggered levels (Figure 1B).

Collectively, these findings robustly affirm the influence of MvfR on mitochondrial functions. These results underscore the efficacy of either the anti-MvfR compound D88 or the SS-31 compound alone in counteracting the MvfR-mediated perturbations observed, thereby averting the disruption of critical mitochondrial functions in the skeletal muscles of infected animals.

### MvfR suppresses the activity of detoxifying enzymes

The increase in ROS production due to MvfR’s contribution to infection led us to investigate the effect of MvfR on the activity of two detoxifying enzymes; superoxide dismutase (SOD) and catalase (CAT). These antioxidant enzymes are essential for maintaining redox homeostasis and protecting cells against oxidative damage (Li et al., 2019). As anticipated, suppression in SOD activity was observed in the skeletal muscle of mice infected with PA14. In contrast, the PA14 isogenic mutant strain Δ*mvfR* exhibited no inhibition of SOD activity and showed levels comparable to those observed in the sham control group or PA14-infected and treated mice with compound D88 or SS-31 (Figure 2A). Furthermore, catalase activity was also reduced in infection with PA14. While infection with the Δ*mvfR* strain resulted in higher catalase activity than in PA14, it was lower than that displayed by the sham control group, indicating that other functions besides MvfR contribute to the decreased catalase activity in PA14-infected mice. Significantly, the administration of SS-31 effectively normalized the levels of catalase activity to levels akin to those observed in the sham control group (Figure 2B). At the same time, D88 reinstated them similarly to that of the Δ*mvfR* strain. Collectively, these outcomes signify that MvfR increased ROS production during infection should also be attributed to suppressed SOD and impaired catalase activity. Notably, the addition of the anti-mvfR compound D88 and the mitochondrial peptide SS-31 successfully counteract MvfR-induced disturbances, underscoring their efficacy in reinstating cellular homeostasis.

**Figure 2:**
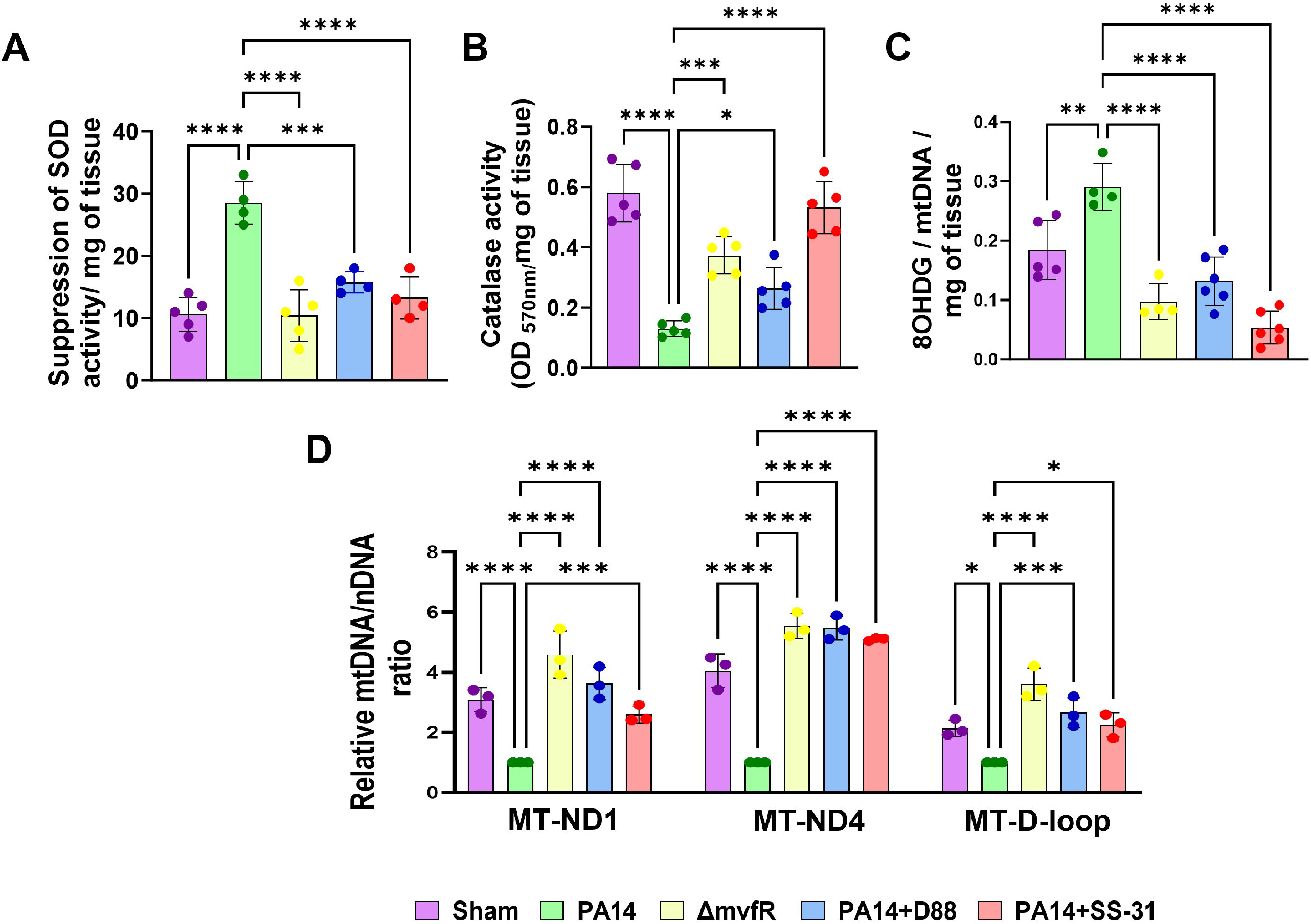
D88 and SS-31 mitigate the MvfR-mediated reduction of the activity of the antioxidant enzymes, and effects on mitochondrial DNA content and oxidative DNA damage. Mice infected with PA14 or Δ*mvfR* strain, or PA14-infected and treated with anti-mvfR compound D88 or mitochondrial-targeted SS-31 peptide were used to assess (A) Suppression of SOD activity represented as percent inhibition (B) Catalase activity at OD_570nm_ (C) Mitochondrial oxidative DNA damage measured by the amount of 8 hydroxy-deoxyguanosine. Gastrocnemius muscle was used to determine (A-C) per milligram of the tissue. Sham represented the baseline control of the burn group. (D) Quantitative real-time PCR (qRT-PCR) analysis of mitochondrially encoded *MT-ND1* gene, *MT-ND4* gene, and *MT-D-loop* genes. Mitochondria were isolated from the gastrocnemius muscle of the above mice groups. Transcript levels were normalized to GAPDH amplified from nuclear DNA. For qPCR analysis, mice infected with PA14 were set as 100% and served as control. Means ± SDs are shown; * p < 0.05, ** p < 0.01, *** p < 0.001, **** p < 0.0001, shows significance. One-way ANOVA followed by Tukey’s post hoc test was applied.

### MvfR promotes oxidative damage of mitochondrial DNA and affects the mitochondrial DNA content, effects that are alleviated by D88 and SS-31

The MvfR-promoted increase in ROS levels prompts us to assess the oxidative damage of mitochondrial DNA by quantifying the 8-hydroxy-deoxyguanosine (8-OH-dG) levels in the gastrocnemius muscle. Compared to the sham control, an increase in oxidative damage of mitochondrial DNA was observed in mitochondria isolated from the skeletal muscle of the PA14-infected mice. Conversely, the PA14 isogenic mutant Δ*mvfR* reduced the levels of 8-OH-dG, opposite to those observed in the infected group (Figure 2C). Furthermore, we explored whether D88 and SS-31 protect mitochondria against oxidative stress-induced damage in PA14-infected mice. Both compounds reversed the phenotype observed in the PA14-infection group (Figure 2C).

To validate the impact of MvfR function on the increase of oxidative damage of mitochondrial DNA, we quantified the mitochondrial encoded DNA (mtDNA) content relative to nuclear-encoded DNA (nDNA) content in the mitochondrial fraction of the gastrocnemius muscle of mice infected with PA14, Δ*mvfR* strain or PA14-infected receiving the additional administration of D88 or SS-31. We determined the mtDNA/nDNA ratio using qPCR to assess the expression of the genes of mitochondrially encoded NADH dehydrogenase subunit 1 (*ND1*), NADH dehydrogenase subunit 4 (*ND4*), and *D-loop*. ND1 and ND4 are involved in mitochondrial electron transport and are localized in the inner mitochondrial membrane. The D-loop region, the main non-coding area of the mtDNA and a hot spot for mtDNA alterations, contains essential transcription and replication elements (46, 47). Compared to *mvfR*-infected mice muscle samples, ∼ 3-fold and 4-fold decrease in mtDNA/nDNA ratio in *ND1* and *ND4* genes was observed in the PA14-infected group, respectively (Figure 2D).

Similarly, a 2-fold reduction in mtDNA/nDNA ratio in the *D-loop* gene was observed in the skeletal muscle of the PA14-infected mice (Figure 2D). Thus, the ratios of mtDNA/nDNA of three mitochondrial genes, *ND1*, *ND4*, and *D-loop*, were significantly decreased due to MvfR’s role in infection, indicating a lower mtDNA content in infected muscle tissues. The observed decrease was restored by administering D88 and SS-31 in the infected animals similarly by 4-fold in the *ND4* gene and 2-3-fold in the *ND1* and *D-loop* gene, reaching levels similar to the baseline control group (Figure 2D). These findings further support the MvfR-induced oxidative damage of mitochondrial DNA, which leads to a significant decrease in the mtDNA content.

### MvfR negatively modulates mitochondrial dynamics

Mitochondrial fission is crucial for several processes, including proper distribution of mitochondria, cytochrome C release, and mitophagy (48). Mitochondrial fission protein 1 (FIS1) is one of the adaptor proteins involved in mitochondrial fission and the recruitment of the dynamin-related protein 1 (DRP1), a cytosolic GTPase protein, to the mitochondrial outer membrane (48–50). We assessed whether MvfR impacts the levels of these proteins using the mitochondrial lysates of the skeletal muscle from the infected mice (Figure 3A). Compared to mitochondrial fraction from muscle tissues of Δ*mvfR-infected* mice, the protein levels of both FIS1 and DRP1 increased in the PA14-infected group. The addition of D88 in the PA14-infected group brought these proteins to about basal levels, while SS-31 decreased the excessive levels of DRP1 but not that of FIS1 (Figure 3A).

**Figure 3:**
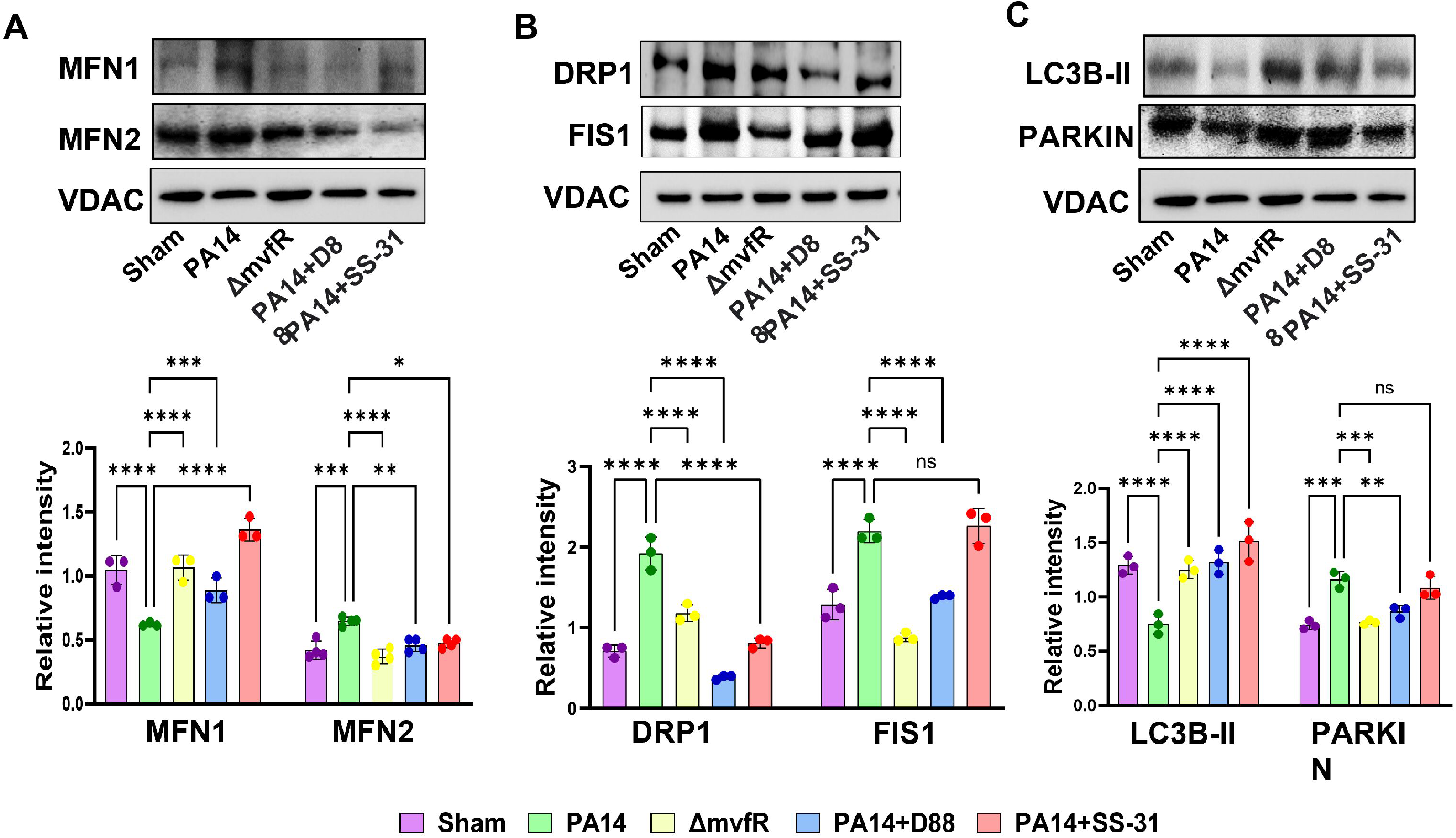
MvfR affects the mitochondrial fusion, fission, and mitophagy-related proteins. These defects can be reinstated by D88 and SS-31. (A) Western blot showing protein levels of mitochondrial shaping proteins MFN1, MFN2 (B) Mitochondrial fission proteins DRP1, FIS1 (C) Mitophagy markers LC3B-II, PARKIN using mitochondrial lysates from the gastrocnemius muscle of mice infected with PA14 or Δ*mvfR* strain, or PA14- infected and treated with anti-mvfR compound D88 or mitochondrial-targeted SS-31 peptide. Sham represents the baseline control of the burn group. (Bottom) Bar charts (bottom) depicting the relative signal intensity versus total protein loading amount for skeletal tissues of three mice (n=3). Error bars represent the standard deviation (SD) from the average of three mice. VDAC was used as a loading control. The statistical difference between each protein level is shown where * p < 0.05, ** p < 0.01, *** p < 0.001, **** p < 0.0001, and ns indicates no significant difference. One-way ANOVA followed by Tukey’s post hoc test was applied.

Moreover, the mitochondrial proteins mitofusin 1 (MFN1) and mitofusin 2 (MFN2) are dynamin-like GTPases essential for regulating the mitochondrial outer membrane fusion dynamics. They are dispensable for mtDNA content, respiratory function, and other critical mitochondrial functions (48). Western blotting analysis showed that in the PA14- infected group, the levels of MFN1 were reduced, and MFN2 slightly increased compared to the protein levels in the muscle tissues of Δ*mvfR*-infected mice. Both D88 and SS-31 reinstated the levels of these proteins in the PA14-infected muscle tissues to similar levels as observed in the Δ*mvfR* or control group, with SS-31 promoting a slightly higher increase in MFN1 (Figure 3B).

Damaged mitochondria are removed by mitophagy (49). Given that our results show that MvfR negatively impacts mitochondrial homeostasis, we investigated whether mitophagy was affected in the skeletal muscle of infected mice. The ubiquitin-mediated mitophagy includes PINK/Parkin. PINK1 activates Parkin and promotes mitophagy through the ubiquitination of mitochondrial proteins. Subsequently, the damaged mitochondria containing ubiquitinated proteins are decorated by autophagosomal microtubule-associated protein 1, light chain 3 isoform B (LC3B), which forms mitophagosomes (Narendra et al., 2010; Eid et al., 2016). We performed western blot analysis of the mitophagy markers LC3B-II and PARKIN using the mitochondrial lysates from the skeletal muscle of infected mice five days post-infection as in all previous experiments. Figure 3C shows that the expression of the ubiquitin E3 ligase, PARKIN, was increased with PA14 infection compared to Δ*mvfR-*infected and sham control groups. Conversely, LC3B-II protein expression was significantly decreased in PA14-infected mice compared to the Δ*mvfR-infected* group, suggesting that MvfR may hinder the clearance of damaged mitochondria. However, adding the anti-*mvfR* compound D88 or SS-31 significantly increased LC3B-II expression in PA14-infected mice, similar to the levels of Δ*mvfR* and sham control (Figure 3C), suggesting that these compounds may help in the elimination of dysfunctional mitochondria in response to PA14 infection. Moreover, in PA14-infected mice receiving D88, PARKIN levels were comparable to Δ*mvfR* and sham control, while SS-31 had no significant effect in restoring PARKIN levels in PA14 infection (Figure 3C). These findings provide additional evidence supporting the impairments in mitochondrial functions mediated by MvfR and the efficacy of D88 in diminishing such impairments.

## DISCUSSION

The importance of the QS transcriptional regulator MvfR and its *mvfR/pqsA-E* system in *P. aeruginosa* virulence and its interrelationship to the other two major QS systems, *lasR/lasI* and *rhrR/rhrI*, has been well established by our group and others (23–26, 37, 51–54). Here, we reveal the impact of this QS transcriptional regulator in the gastrocnemius muscle and provide evidence of its adverse effects on mitochondrial functions in an infection setting *in vivo*. Even though previous studies have implicated MvfR-regulated functions in aspects of mitochondrial dysfunction and MvfR controls many functions that may contribute to the perturbations observed either individually or in synergy, no previous *in vivo* infection studies have addressed the effect of MvfR and its inhibition in skeletal muscle in this setting. This study presents for the first time the consequences of the infection of the key *P. aeruginosa* QS regulator of the *mvfR*/*pqsA-E* system and its regulon inhibition in this setting. This knowledge is pivotal in designing therapeutic approaches that concomitantly inhibit multiple virulence functions, as with the NAM compounds targeting MvfR (37).

Our findings unveiled that five days post-infection, there are multifaceted systemic consequences mediated by MvfR in mitochondrial functions, including bioenergetics, oxidative stress responses, and mitochondrial integrity (Figure 4). Functions that are essential for maintaining mitochondrial homeostasis. ATP production relies on the synchronized function of diverse proteins within the mitochondrial electron transport chain and oxidative phosphorylation (3, 16). Indeed, as the MvfR-mediated perturbations, we found that MvfR reduces ATP production and cytochrome c levels in the murine skeletal muscle, indicating an imbalance in energy homeostasis and impaired bioenergetics. The decrease of cytochrome c protein levels, a pivotal mitochondrial protein involved in electron transport, should contribute to the notably elevated levels of ROS production. Cytochrome c also plays a crucial role in preserving the mitochondrial transmembrane potential necessary for ATP synthesis, corroborating our findings and aligning with studies showing that ATP generated is reduced in the mitochondria of skeletal muscle as a result of alterations in the electron transport chain activity and increased oxidative damage during skeletal muscle aging (55, 56).

**Figure 4:**
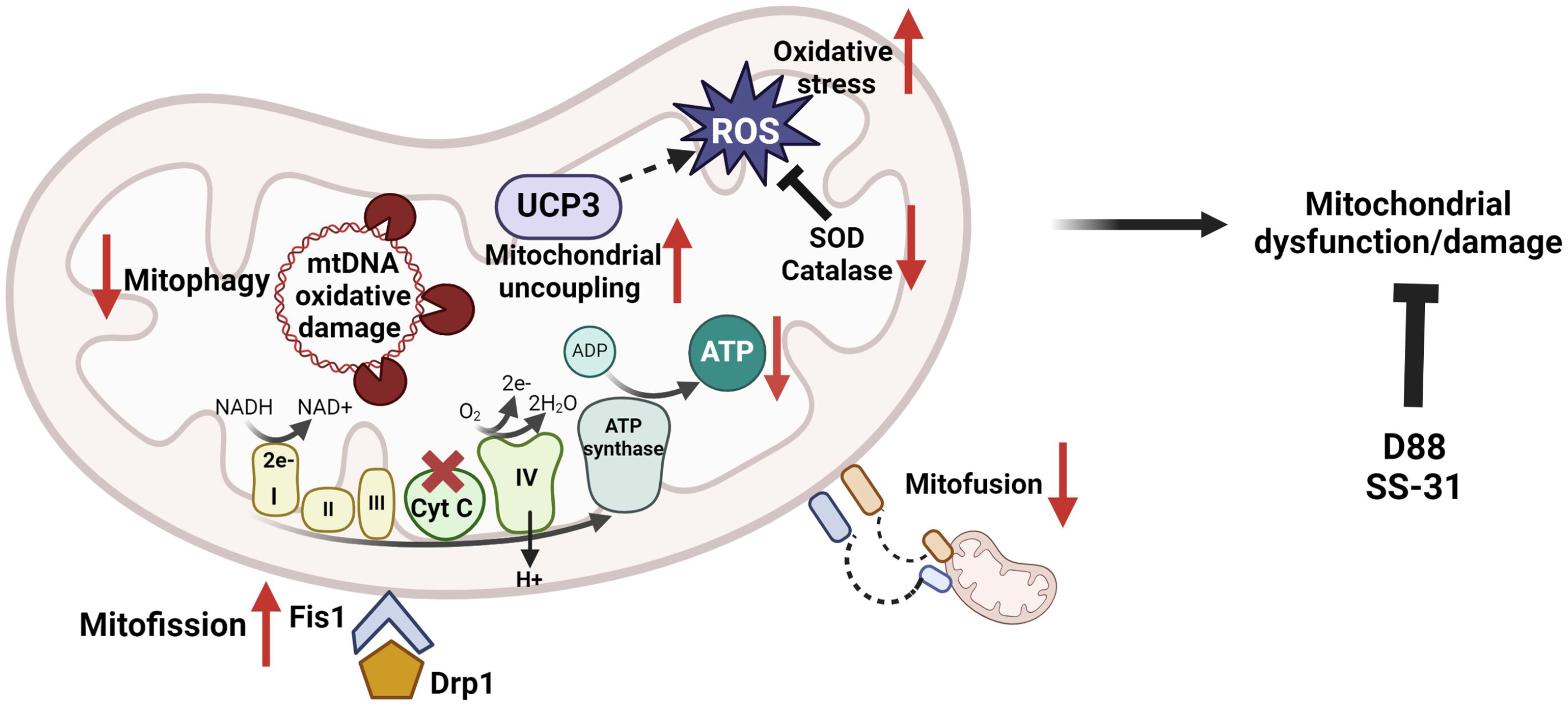
Schematics depicting the perturbations of QS transcriptional regulator MvfR in skeletal murine muscle mitochondria. MvfR decreases ATP levels by reducing expression of the mitochondrial cytochrome c (a component of the electron transport chain), leading to increased ROS production during the oxidative phosphorylation (OXPHOS) process. The decrease in the enzymatic activity of antioxidant enzymes, SOD, and catalase, along with the increase in the mitochondrial uncoupling protein UCP3 contribute to the elevated ROS production. Thus, impaired mitochondrial metabolism and redox deregulation increase ROS levels, causing oxidative damage to mitochondria and a vicious cycle of mitochondrial oxidative stress. These effects result in mitochondrial oxidative DNA damage, which affects the mitochondrial DNA content and impacts mitochondrial quality by altering the levels of fusion and fission proteins. Adding the anti-mvfR compound D88 or the mitochondrial peptide SS-31 rescues the MvfR-mediated mitochondrial defects. Pathways and proteins affected are shown with a red arrow. The figure was created using Biorender.com.

More noteworthy is the elevation of the ROS levels and the accompanied decrease in the activity of the antioxidant enzymes catalase and SOD, which contributes to the elevated levels of ROS observed. ROS can be detrimental to the survival and proliferation of bacteria, raising the question of how *P. aeruginosa* copes with such elevated levels of ROS. These findings emphasize the impact of MvfR on the tightly regulated connections that exist among ATP generation, electron transport chain, and ROS generation, impacting both mitochondrial function and the broader physiological and homeostatic processes within skeletal muscle, offering insights into a mechanism for addressing oxidative stress in the context of *P. aeruginosa* infection. Although modest, the increase in the uncoupling protein UCP3 may also be due to the increase in ROS levels. Although the physiological or pathophysiological role of UCP3 remains unclear and controversial, it is frequently characterized as a regulator of ROS and an uncoupler of mitochondrial oxidative phosphorylation *in vivo* (57).

Another noteworthy impact of the MvfR-mediated increase in ROS is the elevation in oxidative damage of mitochondrial DNA and the perturbation in mitophagy proteins LC3B-II and PARKIN, further instigating mitochondrial dysfunction within murine skeletal muscle. The oxidative mtDNA damage observed may also increase ROS production, leading to mitochondrial dysfunction (58). The mtDNA is essential for the maintenance of normal mitochondrial state and biogenesis. Our results highlight the MvfR-mediated mitochondrial oxidative DNA damage during infection and the decreased mitochondrial DNA content as depicted by the ratio of mtDNA/nDNA in the ND-1 and ND-4 and D-loop region of mitochondria. The increased production of ROS by damaged mitochondria could likely elicit chronic oxidative stress, which may also be responsible for the decrease in the mtDNA content observed. The dysregulation of the mitochondrial dynamics and quality control is also supported by the perturbation of the fusion proteins MFN1 and MFN2 levels, the fission protein DRP1, and its recruiting protein FIS1.

Overall, our data suggest that the QS transcription regulator MvfR changes the protein expression profile of the mitochondrial shaping and mitochondrial division proteins in skeletal murine muscle, contributing to the imbalance of mitochondrial dynamics and impaired mitochondrial function. As shown previously (59) the imbalance in mitochondrial dynamics is characterized by alterations in the fusion processes, leading to reduced ATP production, accumulation of ROS, and oxidative stress. The MvfR-mediated effects underline the role of mitochondrial DNA integrity and its potential impact on skeletal muscle physiology during *P. aeruginosa* infection.

MvfR controls several functions/products that may contribute to the effects observed on mitochondrial dysfunctions. Thus, the findings can also stem from co-actions of more than one of the MvfR-controlled functions (26). For example, phenazines (60, 61), cyanide, the signalling molecules 2-AA and/or HQNO, and PQS although PQS appears not to be relevant in mouse (24, 25, 37). Administration of 2-AA in non-infected mice decreases in skeletal muscle the ATP synthesis rate, compromises muscle contractility, and perturbs the antioxidant defense (30, 34, 36). Recently, in murine infection studies we report mechanistic insights on the role of 2-AA in decreasing bioenergetics and ATP production in macrophages involving the impaired interaction between estrogen-related nuclear receptor alpha (ERRα) and the peroxisome proliferator-activated receptor gamma coactivator 1-alpha (PGC-1α) that impacts the pyruvate transport into mitochondria (62). On the other hand, HQNO, a well-known inhibitor of cytochrome bc1, inhibits the enzymatic activity of mitochondria (63, 64) and, as we showed, also bacterial (65) complex III at the Q_i_. Although the HQNO effect leads to the dysfunction of cellular respiration in *PA,* it is beneficial for *PA* as it promotes biofilm formation and antibiotic tolerance, favoring the persistence of this pathogen in infected tissues (65). More recently, PQS was shown to act as a B-class inhibitor at the I_Q_ site of the mitochondrial complex I (66).

Perhaps the most significant finding of this work is the efficacy of the MvfR NAM inhibitor D88 we developed (37) and the SS-31 mitochondrial-targeted peptide we used to mitigate the mitochondrial derangements that MvfR causes in skeletal muscle. Both compounds restored the derangements observed to almost control levels except for SS-31, which could not restore the levels of the fission protein FIS1, which controls the mitochondrial inner membrane integrity and is dispensable for mitochondrial division. Although SS-31 has been shown to interact with the inner mitochondrial membrane to modulate electron flux, increase ATP generation, and decrease ROS production (67), the alteration in FIS1 due to infection appears not to be rescued by SS-31.

In summary, our study highlights the role of MvfR in orchestrating mitochondrial dysfunction in the gastrocnemius muscle during distant inflicted *P. aeruginosa* infection and underscores the significance of its pharmacological inhibition. Future studies will address mechanistic aspects of the MvfR-promoted mitochondrial dysfunction reported in this study to also aid in developing active therapeutics to prevent the *P. aeruginosa*-mediated mitochondrial dysfunction. The efficacy of the compounds tested to ameliorate the MvfR-mediated aberrations unveils novel avenues for therapeutic interventions and enhanced management against *P. aeruginosa* infections. The prospect of monotherapy or the combination of anti-MvfR agent with antibiotics and/or SS-31 holds promise for optimizing the clinical outcome of patients affected by this highly problematic recalcitrant pathogen.

## MATERIALS AND METHODS

### Bacterial strains and growth conditions

The rifampicin-resistant *P. aeruginosa* human clinical isolate UCBPP-PA14 (PA14) (68) and the PA14 isogenic deletion mutant Δ*mvfR* strain (69) were used in this study. The bacterial strains were grown in lysogeny broth (LB), LB agar plates, or LB agar plates with 100 *μ*g/ml rifampicin. A single colony of PA14 and Δ*mvfR* was inoculated in LB medium, grown at 37°C overnight, and used as a starter culture for an over-day culture by diluting 1:1000 in fresh LB medium. The diluted bacterial culture was further incubated at 37°C until cells reached the optical density (OD) 3.0.

### Pharmacological inhibitors

For all the assays and western blotting, the anti-*mvfR* compound D88 (24mg/kg) (37) or mitochondrial-targeted peptide SS-31 (3mg/kg) (70, 71) was used in the mice infected with the *P. aeruginosa* strain PA14.

### Studies in mice

Animal protocols were approved by the Institutional Animal Care and Use Committee (IACUC) of Massachusetts General Hospital (Protocol no: 2006N000093).

The full-thickness thermal burn injury and infection model (37) was used to determine the role of MvfR, anti-mvfR compound D88, and mitochondrial-targeted peptide SS-31 in skeletal muscle dysregulation by extracting the gastrocnemius muscle of 10-week-old C57BL/6 male mice (Charles River Lab, USA). The mice were randomized into five mice per following groups: sham (burn), infected with PA14 or Δ*mvfR*, or infected with PA14 and treated with D88 or SS-31. The sham group served as baseline control. In the sham group, animals received burn injury and the D88 vehicle (40% captisol in normal saline). Following anesthesia, a thermal burn injury involving 30% total body surface area dorsal burn was produced on the shaved mouse abdomen dermis, and a subcutaneous injection of 500 *μ*l normal saline was administered for spinal protection. An inoculum of ∼3 x 10^5 PA14 or isogenic mutant Δ*mvfR* cells in 100 *μ*l of MgSO4 (10 mM) was injected in the burn eschar intradermally immediately after burn injury. Two groups of mice infected with PA14 also received D88 (24mg/kg) or SS-31 (3mg/kg). For the group supplemented with our MvfR-inhibiting compound D88, mice received subcutaneous injections (at the nape of the animals) starting at 1h and every six hours thereafter till 48 h, and thereafter every 12 h up to 96 h post-infection. Infected mice receiving SS-31 were injected intraperitoneally (IP) starting at 12 h post-infection and then received a second injection at 24 h and thereafter up every 24 h for up to 96 h post-infection. The gastrocnemius muscle was collected after five days post-infection.

### Measurement of oxidative stress

According to the manufacturer’s protocol, the ROS content was measured using an OxiSelect *in vitro* ROS/RNS assay kit (Cell Biolabs Inc., San Diego, CA). Briefly, the gastrocnemius muscle of the sham and mice infected with PA14, Δ*mvfR*, or PA14-infected mice treated with D88 or SS-31 was homogenized in ice-cold PBS. The homogenates were centrifuged at 10,000 rpm for 5 minutes, and the supernatant was used for measuring ROS content. 50 *μ*l of the supernatant and 50 *μ*l of the catalyst was added to each well of a 96-well plate, mixed thoroughly, and incubated for 5 mins at room temperature. Further, 100 *μ*l of fluorescent probe 2,7-dichlorofluorescin diacetate (DCFH-DA) was added to each well, and the plate was incubated for 30 minutes in the dark. The relative fluorescence of the samples and standards was measured using a Tecan plate reader (excitation wavelength of 484 nm and emission wavelength of 530 nm). ROS production was calculated as relative fluorescence units per gram of tissue.

### Measurement of antioxidant activity

The SOD activity of the gastrocnemius muscle tissues from the mice was assessed using the colorimetric assay kit (Abcam, ab65354) as per the manufacturer’s protocol. The muscle tissue was perfused in PBS and homogenized in ice-cold 0.1M Tris-HCl, pH=7.4 containing 0.5% Triton X-100, 5 mM β-mercaptoethanol, 0.1 mg/ml phenylmethylsulphonyl fluorine. The homogenized muscle tissue was centrifuged at 14,000 X g for 5 mins at 4°C, and the supernatant was collected. 20 *μ*l of supernatant, controls, and standards were added in different wells of a 96-well microtiter plate, and 200 *μ*l of water-soluble tetrazolium salt (WST) and 20 *μ*l of enzyme working solution was added to each well. The plate was incubated at 37°C for 20 mins at room temperature. The plate was read in a Tecan microplate reader at 450 nm. The reduction in WST-1 is inhibited by SOD which leads to the dismutation of superoxide radicals to generate hydrogen peroxide and oxygen. Thus, SOD activity was calculated based on the percent inhibition of WST-1 reduction, which was equivalent to the percent inhibition of the superoxide anions.

The catalase activity of the muscle tissues was determined using the catalase activity kit (Abcam, ab83464) according to the manufacturer’s protocol. Briefly, the tissue was homogenized in 200 *μ*l of ice-cold assay buffer and centrifuged at 14,000 rpm for 15 mins at 4°C. The supernatant was used for the further assay as described in the protocol. The 96-well plate containing the sample, standards, and controls with the catalase reaction mix in different wells was incubated at 25°C for 30 mins. The catalase activity was determined by measuring the optical density at 570 nm in a Tecan microplate reader.

### Measurement of ATP

The amount of ATP generated in the gastrocnemius muscle samples was measured using the ATP assay kit (Abcam, 83355) described in the manufacturer’s protocol. The muscle tissues were washed in ice-cold 1X PBS, homogenized in 100 *μ*l ice-cold 2N perchloric acid, and kept on ice for 30-45 mins. The tissue samples were centrifuged at 13,000 X g for 2 mins at 4°C. The supernatant was collected and used for the assay described in the protocol. After 30 minutes of incubation, the reaction was stopped, and the optical density was measured at 570 nm in a Tecan microplate reader. The amount of ATP generated was calculated as optical density at 570 nm per gram of tissue.

### Mitochondria isolation

The gastrocnemius muscle tissue was suspended in 2 ml of ice-cold mitochondrial isolation buffer (Abcam, ab110168) with a 25X protease inhibitor cocktail (Millipore 1183580001) and was kept on ice for 1 h. The muscle tissue was homogenized using a Dounce homogenizer at 4°C. The homogenized sample was centrifuged at 2,000 rpm for 5 mins at 4°C to isolate the nuclear fraction. The supernatant was further centrifuged at 12000 rpm for 10 mins to obtain a pellet enriched with mitochondria. The supernatant was discarded, and the mitochondrial pellet was lysed in 200 *μl* of lysis buffer. The mitochondrial lysate was stored at −80°C until further use.

### Oxidative damage of Mitochondrial DNA (mtDNA)

Mitochondrial DNA damage was assessed in the mitochondrial fraction by the content of 8-hydroxydeoxyguanosine (8-OHdG) according to the manufacturer’s protocol (STA-320, Cell Biolabs). Briefly, the 96-well microplate was coated with 100 *μl* of 8-OHdG conjugate (1 *μg*/ml in 1x PBS) and incubated overnight at 4°C. The next day, the 8-OHdG coating solution was removed by washing with distilled water. Blocking was carried out with 200 *μl* of the assay diluent for 1 h at room temperature. Subsequently, the plate was transferred to 4°C until further use. Simultaneously, DNA was isolated from the mitochondrial lysate obtained from the gastrocnemius muscle tissue of the mice using the DNeasy Blood and Tissue kit (Qiagen). 1 *μg*/*μl* of the extracted DNA was converted to single-stranded DNA by incubating the sample at 95°C for 5 mins. The DNA samples were digested to nucleosides by incubating with 10 units of nuclease P1 buffer for 2 h at 37°C. Next, 5 units of alkaline phosphatase with 100 mM Tris buffer, pH 7.5 was added, and the tubes were incubated for 1 h at 37°C. The reaction mixture was centrifuged for 5 mins at 6000 X g, and the supernatant was collected for use in ELISA. The assay diluent was removed from the 8-OHdG conjugate-coated plate, and 50 *μl* of the sample was added. The plate was incubated for 10 mins on an orbital shaker. After 10 mins incubation, 50 *μl* of anti-8-OHdG antibody was added, and the plate was further incubated for 1 h at room temperature on an orbital shaker. After 60 mins, the plate was washed three times with 250 *μl* of 1X wash buffer, followed by adding 100 *μl* of the secondary antibody-enzyme conjugate and incubating for 1 h on an orbital shaker. After 60 mins incubation, the plate was rewashed with 1X wash buffer, and 100 *μl* of the substrate solution was added. The plate was further incubated for 30 minutes at room temperature in an orbital shaker, and the enzymatic reaction was stopped immediately by adding 100 *μl* of the stop solution. Absorbance was read at 450 nm in a Tecan microplate reader.

### mtDNA Quantification

The mtDNA present per nuclear DNA in the mitochondrial fraction of the gastrocnemius muscle was quantified by quantitative PCR using the following primers: mitochondrial ND1 forward primer, CCTATCACCCTTGCCATCAT; mitochondrial ND1 reverse primer, GAGGCTGTTGCTTGTGTGAC; mitochondrial ND4 forward primer, AACGGATCCACAGCCGTA; mitochondrial ND4 reverse primer, AGTCCTCGGGCCATGATT; and mitochondrial D-loop forward primer, AATCTACCATCCTCCGTGAAACC mitochondrial D-loop reverse primer, TCAGTTTAGCTACCCCCAAGTTTAA. The GAPDH gene was used to quantify the nuclear DNA from the nuclear fraction using the GAPDH forward primer, AGGCCGGTGCTGAGTATGTC, and the GAPDH reverse primer, TGCCTGCTTCACCACCTTCT. The relative mitochondrial content was quantified using the difference between mtDNA and nuclear DNA by ΔΔC(t) method (72, 73).

### Western Blotting

As described above, the mitochondrial lysates were prepared in RIPA lysis buffer (Cell Signalling Technology, USA). The lysate concentration was determined from each sample by a bicinchoninic acid (BCA) protein assay kit (Thermo Fisher Scientific, USA). 40 -50µg of mitochondrial lysate were prepared in RIPA buffer with 1X Laemmili buffer, boiled for 10 mins at 95°C, and stored at −80°C. The mitochondrial fractions were separated by SDS PAGE and transferred to the PVDF membrane (Bio-Rad). After blocking with 2.5% BSA in TBS containing 0.1% Tween 20 for 1 h at room temperature, the membranes of the mitochondrial lysates were incubated overnight with the primary antibodies specific for cytochrome c (Cell Signalling Technology, 136F3), UCP3 (Cell Signalling Technology, D6J8K), LC3B-II (Cell Signalling Technology, 2775S), VDAC (Cell Signalling Technology, D73DI2), PARKIN (sc-32282), MFN1 (ab126575), MFN2 (ab56889), DRP1 (AB184247) and FIS1 (AB229969). After washing, the membranes were incubated with anti-rabbit secondary antibody for cytochrome c, UCP3, LC3B-II, VDAC, MFN1, DRP1, FIS1, and anti-mouse antibody for PARKIN, MFN2, and β-actin. The bands were detected by Supersignal West Pico Chemiluminescent substrate (Thermo Scientific) reaction, and the membrane blots were visualized in the ChemiDoc imaging system (Bio-rad Laboratories). The densitometric analysis of the bands was done using ImageLab software.

### Statistical analysis

For the assessment of ATP, ROS, catalase, SOD, and mtDNA damage, gastrocnemius muscle tissues from five different mice were used individually (n = 5). For mtDNA content and western blotting, mitochondria were isolated from muscle tissues of mice n = 3 and used individually. The data were analyzed using one-way variance (ANOVA) followed by Tukey’s post hoc t-test and plotted using GraphPad Prism software. For all experiments, P<0.0001 was considered significant.

## ACKNOWLEDGEMENTS

This work was supported by the NIH award R01AI134857, The John Lawrence Massachusetts General Hospital Research Scholar Award and Shriner’s grant 83009 to L.G.R., and the Shriner’s grant 85132 to A.A.T. The funders had no role in the study design, data collection, analysis, publication decision, or manuscript preparation.

**L**.G.R. has a financial interest in Spero Therapeutics, a company developing therapies to treat bacterial infections. L.G.R.’s financial interests are reviewed and managed by Massachusetts General Hospital and Partners Health Care in accordance with their conflict-of-interest policies. No funding was received from Spero Therapeutics, and it had no role in study design, data collection, analysis, interpretation, or the decision to submit the work for publication. The remaining authors declare no competing interests.

## Authors Contributions

L.G.R. and A.A.T. conceptualization, S.A., V.K.S., A.C. A.A.T. and L.G.R., designed research; S.A., V.K.S., S.C., A.D., C.dC., O.I., L.Y., R.B., performed research; S.A., V.K.S., A.C., and L.G.R. analyzed data; and S.A., A.D., and L.G.R. wrote the paper.

